# Transcriptomic analysis reveals immune signatures associated with specific cutaneous manifestations of lupus in systemic lupus erythematosus

**DOI:** 10.1101/2025.04.27.649460

**Authors:** Ernest Y. Lee, Sarah Patterson, Zachary Cutts, Cristina M. Lanata, Maria Dall’Era, Jinoos Yazdany, Lindsey A. Criswell, Anna Haemel, Patricia Katz, Chun Jimmie Ye, Charles Langelier, Marina Sirota

**Affiliations:** Department of Dermatology, University of California, San Francisco, San Francisco, CA, United States; Division of Rheumatology, Department of Medicine, University of California, San Francisco, San Francisco, CA, United States; Division of Infectious Diseases, Department of Medicine, University of California, San Francisco, San Francisco, CA, United States; Department of Pediatrics, University of California, San Francisco, San Francisco, CA, United States; Bakar Computational Health Sciences Institute, University of California, San Francisco, San Francisco, CA, United States; Genomics of Autoimmune Rheumatic Disease Section, National Human Genome Research Institute, National Institutes of Health, Bethesda, MD, United States

**Author notes:** Corresponding Authors: Marina Sirota Bakar Computational Health Sciences Institute, University of California, San Francisco, San Francisco, CA, United States.

## Abstract

Systemic lupus erythematosus (SLE) presents with diverse heterogenous cutaneous manifestations. However, the molecular and immunologic pathways driving specific cutaneous manifestations of SLE are poorly understood. Here, we leverage transcriptomics from a large well-phenotyped longitudinal cohort of SLE patients to map molecular pathways linked to ten distinct SLE-related rashes. Through whole blood and immune cell-sorted bulk RNA sequencing, we identified immune signatures specific to cutaneous subtypes of SLE. Subacute cutaneous lupus (SCLE) exhibited broad upregulation of interferon, TNF-α, and IL6-JAK-STAT3 pathways suggesting potential unique therapeutic responses to JAK and type I interferon inhibition. While interferon signaling is prominent in SCLE, discoid lupus, and acute lupus, it is unexpectedly attenuated in patients with skin and mucosal ulcers. Pathway and cell-type enrichment analysis revealed unexpected roles for CD14+ monocytes in photosensitivity of SLE and NK cells in alopecia, mucosal ulceration, and livedo reticularis. These findings illuminate the immune heterogeneity of rashes in SLE, highlighting subtype-specific mechanistic targets, and presenting opportunities for precision therapies for SLE-associated skin phenotypes.

## Introduction

Systemic lupus erythematosus (SLE) is a chronic autoimmune inflammatory disorder characterized by broad and heterogenous clinical manifestations that reflect complex, dysregulated immune responses (1). Cutaneous lupus erythematosus (CLE) encompasses various mucocutaneous manifestations, including acute cutaneous lupus (ACLE, includes malar rash), chronic cutaneous lupus erythematosus (CCLE, includes discoid lupus and chilblain lupus), subacute cutaneous lupus erythematosus (SCLE), and lupus panniculitis. However, lupus also manifests clinically as alopecia, skin and mucosal ulcers, and livedo reticularis (2, 3). A large subset of patients with SLE will exhibit cutaneous lupus, though not all patients with cutaneous lupus will progress to SLE(4). These distinct cutaneous manifestations not only contribute to disease burden but also hint at underlying immunological heterogeneity, suggesting that different pathways may drive specific phenotypes. Despite advances in our understanding of SLE immunopathology, the immune drivers of specific SLE-related cutaneous manifestations amongst patients with SLE remain poorly understood.

At present, we are unable to predict or identify which patients with SLE will exhibit specific cutaneous manifestations. Although a subset of SLE patients will initially present with a rash, many patients may initially present with lupus nephritis or neurologic symptoms. Although malar rash and discoid rash are part of the classification criteria for SLE, we now recognize that there are other cutaneous manifestations of SLE, not part of the classification criteria, that have significant clinical and psychosocial impacts on SLE patients (2, 5). Since SLE is an immunologically complex disorder, identifying molecular pathways that are associated with specific SLE-related cutaneous lesions and illuminating new molecular targets is of utmost clinical importance.

High-throughput transcriptomic profiling has proven valuable in mapping immune dysregulation in autoimmune diseases and identifying disease-specific clinically relevant gene signatures across many autoimmune and inflammatory conditions including SLE (6–11). To study the SLE disease heterogeneity, identify biomarkers and discover therapeutic targets, several studies have utilized bulk RNA sequencing (bulk RNAseq) of whole blood (8), as well as single-cell RNA sequencing (scRNAseq) of peripheral blood mononuclear cells (PBMCs) (12) to identify specific gene signatures in cell types crucial to lupus pathogenesis. In addition, they have identified key gene signatures related to type I interferon signaling in B cells, plasma cells, and granulocytes in SLE (13–15). Prior work also has showed how the American College of Rheumatology (ACR) classification criteria of SLE clusters into subgroups that are related to specific differences in clinical phenotypes and lupus severity (16)(17), and identified race-specific and cell-type specific immunologic differences in clinical clusters of SLE patients (18). Transcriptomic analysis previously identified differentially expressed transposable elements across all phenotypes that were cell-type and phenotype specific, and were related to genes primarily involved in antiviral signaling and IFN signaling (19). Methylation analysis of SLE patients identified 256 differentially methylated CpGs across disease associated with ethnic-associated clinical outcomes (16). Several prior studies have also utilized bulk RNAseq and scRNAseq on skin biopsies from healthy patients and SLE patients with discoid lupus or acute cutaneous lupus to identify a role for infiltration of plasmacytoid dendritic cells and upregulation of an associated type I interferon signature in cutaneous lesions (20–23). These previous studies focused primarily on differentiating SLE patients from healthy controls and identifying gene signatures in whole blood related to global disease activity and cytokine profiles, and clinical manifestations such as lupus nephritis. However, few studies have focused on identifying immunologic pathways in the blood that are predictive of cutaneous phenotypes, and none have a large enough sample size to encompass the entire clinical spectrum of SLE-related cutaneous manifestations. Furthermore, these studies are often limited to smaller cohorts with more homogeneous and less diverse demographics, limiting the predictive power and generalizability.

Leveraging a large, high-quality ethnically diverse longitudinal cohort of >400 SLE patients with carefully curated clinical data from the California Lupus Epidemiology Study (CLUES), we identify molecular correlates associated with specific cutaneous features of lupus. Through parallel whole blood and immune cell-sorted bulk RNA sequencing, we describe transcriptional profiles across ten distinct subtypes of SLE-related rashes, enabling us to characterize immune pathways and cellular contributors with unprecedented specificity. Our findings reveal a diverse landscape of immune signatures underlying subtypes of SLE-related cutaneous manifestations.

## Results

### Transcriptomic landscape of cutaneous manifestations of SLE in the CLUES cohort

Participants were recruited from the California Lupus Epidemiology Study (CLUES). CLUES is a multi-racial/ethnic cohort of patients with confirmed SLE (Figure 1). SLE diagnosis was confirmed by board certified study physicians based on the ACR revised criteria from 1982 and updated in 1997 or based upon a confirmed diagnosis of lupus nephritis. These patients were longitudinally followed from time of enrollment with careful annotation of clinical characteristics and molecular measurements (Figure S1). In this study, we focused on 10 different rashes and/or cutaneous manifestations related to SLE: acute cutaneous lupus (malar rash), discoid lupus (DLE), photosensitivity, mucosal ulcers, skin ulcers, alopecia, subacute cutaneous lupus (SCLE), lupus panniculitis, livedo reticularis, and other rashes related to SLE (Figure 1). We excluded other clinical phenotypes due to low incidence. We generated two datasets to be analyzed together. Bulk RNA sequencing of whole blood was conducted on N = 461 samples taken from 276 CLUES patients. The CLUES cohort represents both female and male patients, though is majority female, across eight different self-reported races spanning a range of low to high lupus severity as measured by standardized clinical criteria (lupus severity index) (Table 1). We also evaluated a subset of N = 120 samples from SLE patients with cell-sorted bulk RNA sequencing of CD4+ T cells, CD14+ monocytes, B cells, and NK cells (Figure 1).

**Figure 1:**
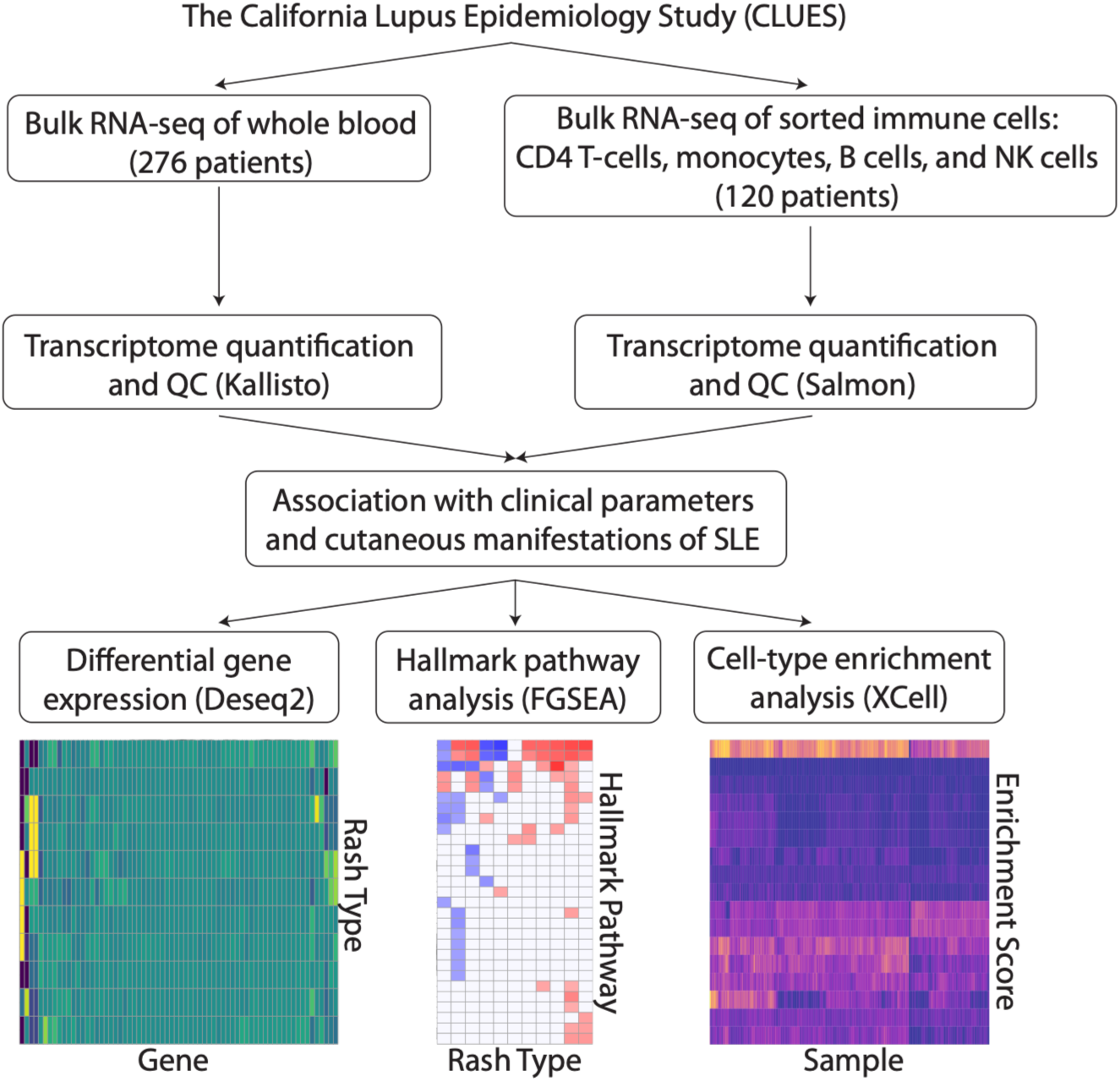
Study overview and design. Patients were recruited from the CLUES cohort and bulk RNA sequencing was conducted. Data was processed for transcriptome quantification and quality control. We then associated clinical parameters with cutaneous manifestations of SLE, and downstream conducted differential gene expression analysis, pathway analysis, and cell type enrichment analysis.

**Table 1:** CLUES cohort demographic characteristics.

| Characteristics | Overall (N = 276) |
| --- | --- |
| Sex |  |
| Female | 244 (88.4%) |
| Male | 32 (11.6%) |
| Age | 44.5 ± 14.0 years |
| Race |  |
| White | 99 |
| Black | 29 |
| Asian | 102 |
| American Indian | 1 |
| Mixed | 10 |
| Other | 32 |
| Unknown race | 3 |
| Lupus severity index | 6.9 ± 1.6 |
| Malar rash | 139 (50.0%) |
| Discoid rash | 40 (14.4%) |
| Photosensitivity | 103 (37.3%) |
| Mucosal ulcers | 120 (43.5%) |
| Alopecia | 110 (39.9%) |
| Panniculitis | 9 (3.3%) |
| SCLE | 18 (6.5%) |
| Skin Ulcers | 12 (4.3%) |
| Livedo reticularis | 32 (11.6%) |
| Other SLE related rash | 35 (12.7%) |
| Other Non-SLE rash | 25 (9.1%) |

After quality control (Figure S2) and comprehensive analysis of the clinical metadata to identify specific dermatologic manifestations of lupus, we then conducted differential gene expression, pathway analysis, and cell-type enrichment on each clinical phenotype to identify transcriptomic and immunologic signatures associated with SLE related rashes (Figure 1). The CLUES cohort represents both female and male patients, though is majority female, across eight different self-reported races spanning a range of low to high lupus severity as measured by standardized clinical criteria (SLE disease activity index (SLEDAI) score and lupus severity index). Most patients in this cohort had a history of malar rash, mucosal ulcers, photosensitivity, and alopecia, with a smaller subset of patients with DLE or SCLE (Figure 2A). Lupus panniculitis, skin ulcers, livedo reticularis were less common. Principal component analysis (PCA) plots of the overall bulk transcriptomic data from whole blood for subpopulations of patients with each of the different SLE-related rashes are shown (Figure 2B, Figure S3). In the first two principal components, there is no clear separation or clustering of the clinical subsets of the patients with or without a specific SLE-related rash, suggesting that more subtle molecular effects may be responsible for different clinical findings. No significant batch effect is seen with the two runs of samples being collected in parallel. All annotated phenotypes of SLE-related cutaneous manifestations are shown together with the raw dataset of bulk RNAseq counts across all samples (Figure S4).

**Figure 2:**
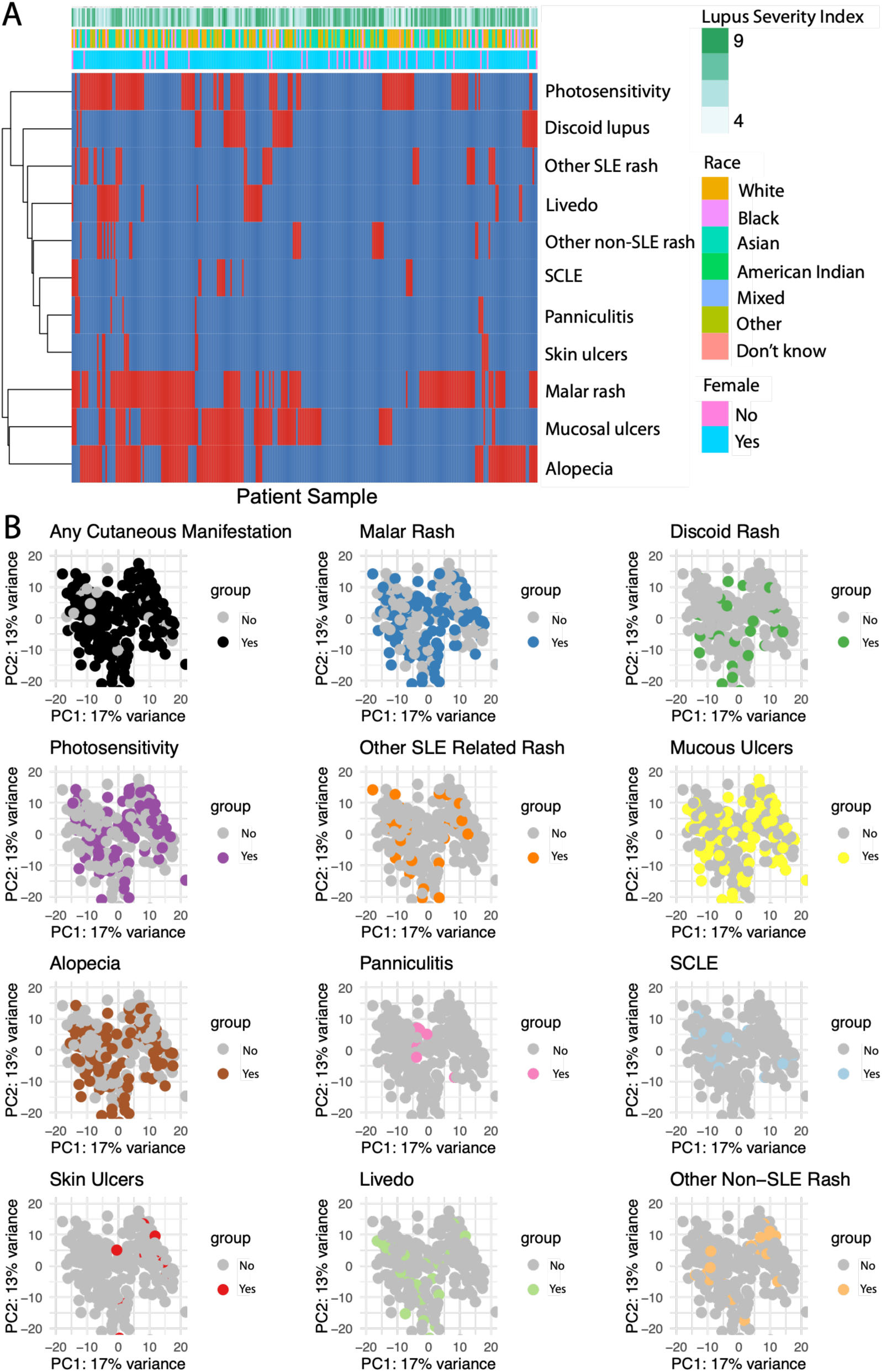
Transcriptomic landscape of cutaneous manifestations of SLE in the CLUES cohort. (A) Clinical characteristics and cutaneous subtypes of lupus across the entire cohort. (B) PCA plots showing whole blood bulk RNAseq and each associated SLE-related rash subtype.

### Interactome and disease signatures of distinct SLE-related rashes

To further identify molecular differences underlying SLE-related rashes, we conducted differential gene expression analyses with Deseq2 (24). First, we conducted differential expression on the whole blood transcriptomes for each clinical phenotype while controlling for demographic features and clinical parameters such as sex, race, medication use, replicates, and batch, and then compared our findings to the differential gene expression on the cell-sorted bulk transcriptomes from CD4 T cells, CD14 monocytes, CD19 B cells, and NK cells. For each differential gene expression analysis, we compared patients with and without specific cutaneous manifestations of lupus. There were 2 differentially expressed genes (DEGs) (FDR < 0.05) in the blood of patients with discoid lupus, 6 DEGs in patients with alopecia, 4 DEGs in SCLE, 1 DEGs in malar rash, 2 DEGs in livedo, 6 DEG in skin ulcers, 4 DEGs for mucosal ulcers, and 2 DEGs for photosensitivity (Table 2). 8 DEGs were identified for lupus panniculitis. Utilizing the identities of these genes, we were able to generate an interactome of SLE-related rashes based on shared DEGs between each pairwise combination of cutaneous phenotypes (Figure 3A, Figure S5A). We then calculated the log-2-fold-change (log2FC) for the union of each of the identified DEGs (34 total unique genes) from all clinical phenotypes across whole blood to visualize the differential expression fingerprint for each gene. Hierarchical clustering of these manifestations shows similarities between the expression fingerprints of lupus panniculitis, skin ulcers, and SCLE (Figure 3A).

**Figure 3:**
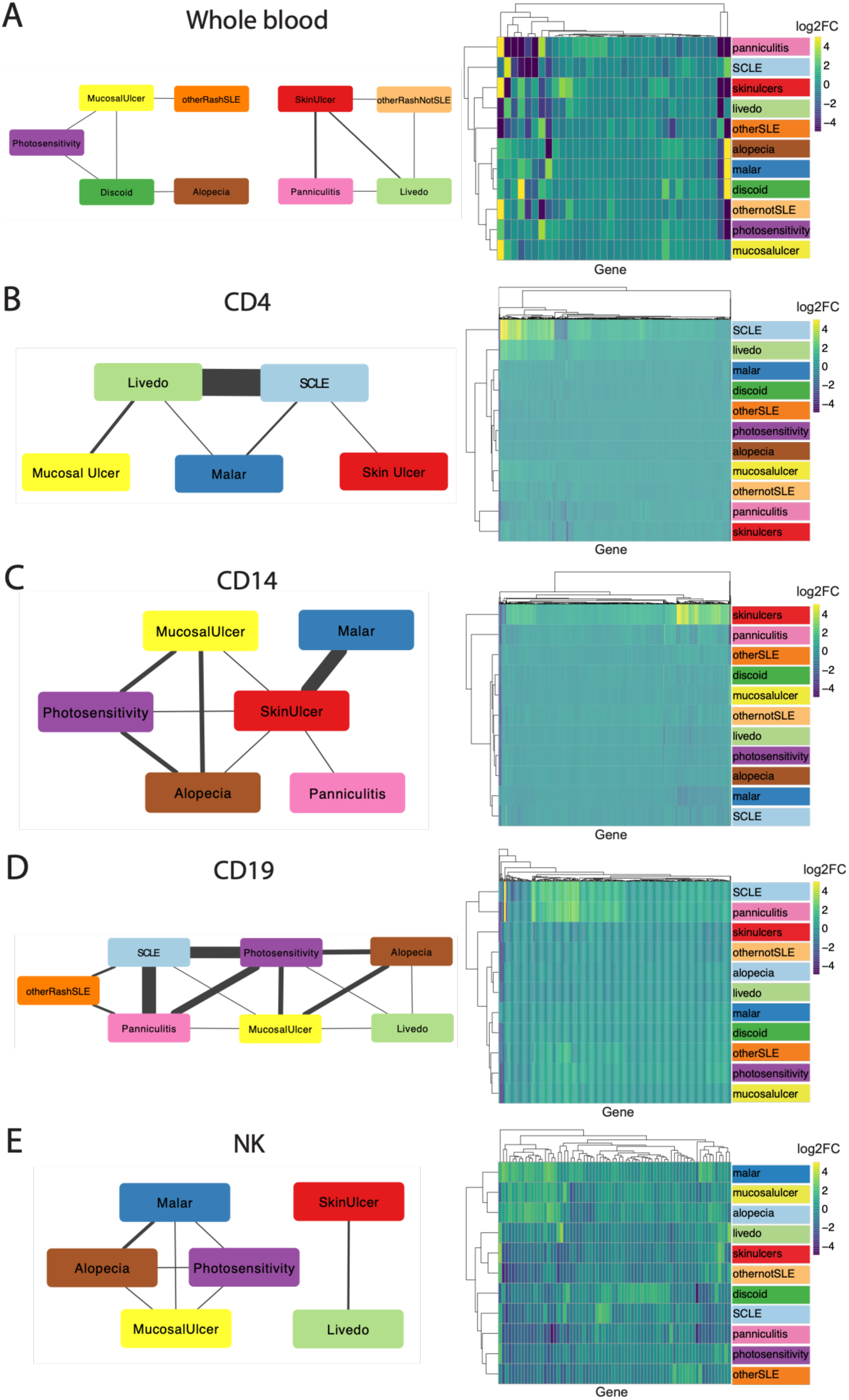
Interactome and disease signatures of distinct SLE-related rashes. Differential gene expression analysis identified interactome and disease signatures of each distinct SLE rash subtype, shown for whole blood (A), and in immune cell-sorted data for CD4 T cells (B), CD14 monocytes (C), CD19 B cells (D), and NK cells (E). Networks of each cutaneous manifestation is constructed based on number of differentially expressed genes in common. Edge thickness is directly proportional to the number of overlapping differentially expressed genes in common between clinical manifestations. Heatmaps show upregulated (yellow) and downregulated (blue) differentially expressed genes by log2-fold-change taken from the union of all differentially expressed genes across cutaneous phenotypes.

**Table 2:** Differentially expressed genes by SLE cutaneous phenotype type in whole blood and immune cell sorted data.

| Clinical Feature | Whole blood | CD4 (T cell) | CD14 (monocyte) | CD19 (B cell) | NK cell |
| --- | --- | --- | --- | --- | --- |
| Malar | 1 | 58 | 16 | 0 | 20 |
| Discoid | 2 | 1 | 1 | 0 | 18 |
| Photosensitivity | 2 | 1 | 6 | 198 | 5 |
| Other rash SLE | 6 | 1 | 0 | 4 | 6 |
| Mucosal ulcers | 4 | 5 | 7 | 16 | 5 |
| Alopecia | 6 | 0 | 7 | 7 | 19 |
| Panniculitis | 8 | 2 | 6 | 85 | 0 |
| SCLE | 4 | 737 | 0 | 178 | 4 |
| Skin ulcers | 6 | 5 | 572 | 2 | 2 |
| Livedo | 2 | 258 | 1 | 1 | 5 |
| Other rash | 2 | 0 | 0 | 0 | 0 |

Since comparatively fewer differentially expressed genes were identified from the whole blood RNAseq, we then conducted differential gene expression on each SLE-related rash for the bulk transcriptomes from sorted immune cells from a subset of the CLUES cohort. We looked at differential gene expression in CD4 T cells, CD14 monocytes, CD19 B cells, and NK cells, to identify cell-type specific gene signatures for each cutaneous clinical phenotype of interest. We then plotted the log2FC for the union of all DEGs for each cell type across phenotypes. The expression fingerprints of all DEGs in the CD4 T-cells (821 total genes) were plotted for all clinical phenotypes (Figure 3B). In CD4 T-cells, livedo reticularis and SCLE shared 238 DEGs, while 3 DEGs were shared between livedo and mucosal ulcers, 2 DEGs shared between malar rash and SCLE, and 1 gene shared between SCLE and skin ulcers (Figure 3B, Figure S5B). Most of the DEGs identified in the CD4 cohort were upregulated genes. Hierarchical clustering of the clinical phenotypes showed similarity between patients with skin ulcers and lupus panniculitis, while SCLE and livedo were also closely similar, given the large number of shared DEGs.

In the CD14 monocytes, surprisingly we identified 14 common DEGs between the malar rash and skin ulcer phenotypes, and less than 4 shared DEGs between patients with photosensitivity, mucosal ulcers, and alopecia (Figure 3C, Figure S5C). Expression fingerprints of all DEGs (589 total genes) in CD14 cells showed closer similarity between patients with skin ulcers and panniculitis, but opposite gene expression patterns between malar rash and skin ulcers (Figure 3C). In the CD19 B cells, the gene expression fingerprints of all DEGs in the CD19 cells (383 total genes) showed significant similarity between SCLE and panniculitis with hierarchical clustering. The most DEGs in common were identified between SCLE and panniculitis, SCLE and photosensitivity, and panniculitis and photosensitivity (Figure 3D, Figure S5D). There were 47 DEGs shared between SCLE and panniculitis, 17 shared between SCLE and photosensitivity. Finally, in the NK cells, the clinical phenotypes clustered separately based on shared DEGs. Fewer total DEGs were seen across all phenotypes (72 total genes) compared to the CD4, CD14, and CD19 cell types (Figure 3E, Figure S5E). Very few DEGs were shared between clinical phenotypes, compared to larger overlaps in the other cell types. Only 3 shared DEGs were identified for alopecia and malar rash, 2 DEGs for skin ulcers and livedo, and 1 DEG for other phenotypes.

### Global and cell-type specific immunologic pathways dysregulated in cutaneous phenotypes of SLE

Next, we asked which molecular pathways are dysregulated in the whole blood of patients with cutaneous manifestations of lupus. We calculated the differential gene expression profiles from Deseq2 for each cutaneous phenotype and conducted gene set enrichment analysis using FGSEA (25). We utilized the Hallmark pathways from MSigDB (26) for each SLE-related rash. We identified several significant upregulated and downregulated pathways for each SLE-related rash in whole blood (padj < 0.05) (Figure 4A, Figure S6).

**Figure 4:**
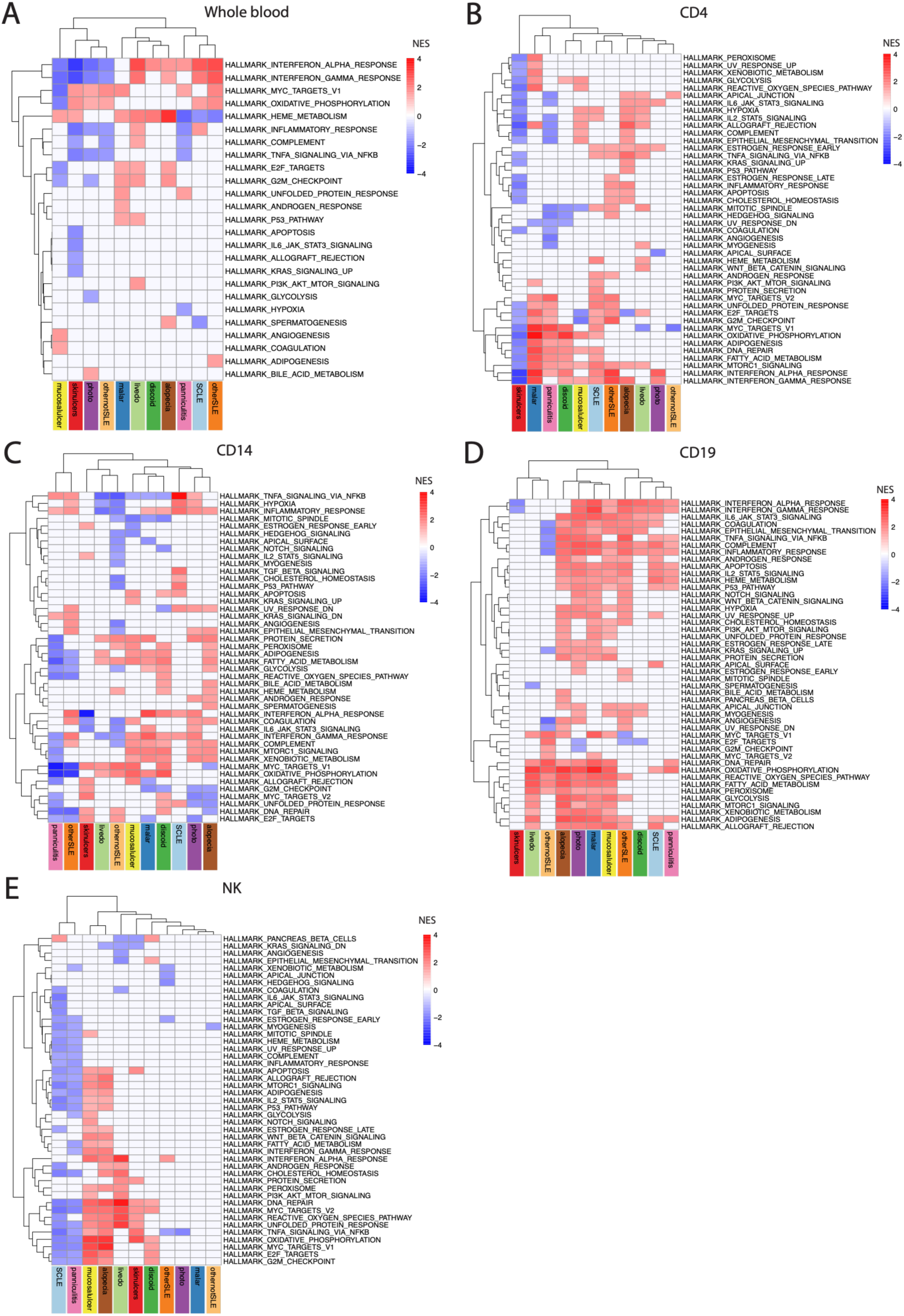
Unsupervised clustering of signaling pathways defining cutaneous manifestations of SLE. Unsupervised hierarchical clustering of normalized enrichment scores for biological pathways from GSEA of bulk RNAseq data from whole blood (A), CD4+ T cells (B), CD14+ monocytes (C), CD19+ B cells (D), or NK cells (E). Only significant (FDR < 0.05) upregulated (red) or downregulated (blue) pathways are shown.

In whole blood, interferon alpha and interferon gamma response were globally upregulated across patients with discoid rash, alopecia, SCLE, livedo, and lupus panniculitis (Figure 4A). Surprisingly, interferon alpha and gamma pathways were downregulated in patients with skin ulcers and mucosal ulcers, potentially suggesting that type I interferon does not directly drive these clinical sub-phenotypes. From hierarchical clustering, SCLE and lupus panniculitis clustered together in terms of pathway signatures. We also observed that the pathway signatures more closely clustered for malar rash, discoid rash, livedo, and alopecia. Separately, we identified that activation of the heme metabolism pathway was the strongest in patients with alopecia. Lupus panniculitis exhibited upregulation of interferon response pathways, it exhibits downregulation of heme metabolism, TNF signaling, and UV response pathways.

Now that we identified global pathway signatures for each clinical phenotypes, we asked whether pathways detected in specific SLE-related rashes were specific to certain cell types. More specifically, we carried out similar differential gene expression analyses for CD4, CD14, CD19, and NK cells (Figure 4B-E). CD4 T cells exhibited upregulation in IFN alpha and gamma related pathways across several phenotypes including malar rash, discoid rash, SCLE, panniculitis, and photosensitivity (Figure 4B). The mammalian target of rapamycin (mTOR) pathway which is relevant to CD4 T cell activation and proliferation was also upregulated. Numerous pathways related to metabolism and cellular respiration and turnover was also upregulated. Interestingly, TNF pathway activation was only seen in a smaller subset of phenotypes, including SCLE, alopecia, and livedo reticularis, but not in malar rash or discoid lupus. The JAK/STAT pathways related to IL2 and IL6 were more strongly prominent in livedo, alopecia, and photosensitivity, which clustered together with hierarchical clustering.

CD14 monocytes and macrophages are known to be dysregulated in SLE, but their role in development of cutaneous lupus is poorly understood. Previous studies have shown that monocytes in SLE exhibit defective phagocytosis as well as aberrant activation in disease initiation and mediation of end organ damage (27). Globally, CD14 monocytes exhibit varied differential pathway expression across cutaneous phenotypes. Interferon alpha and gamma pathways were upregulated in malar rash, discoid rash, SCLE, and photosensitivity. Across all pathways, malar and discoid rash clustered together along with mucosal ulcers, while SCLE, photosensitivity, and alopecia clustered together (Figure 4C). Surprisingly, TNF signaling was upregulated in SCLE, lupus panniculitis, and photosensitivity, but downregulated across malar and discoid rash, and mucosal ulcers. CD14 monocytes also exhibited significant expression of downregulated UV response pathways in SCLE and photosensitivity, suggesting that they are involved in regulating immune responses to UV radiation in those conditions.

CD19 B cells exhibited the broadest number of upregulated proinflammatory pathways of all the cell types studied. It is well known that B cells play a central role in the pathogenesis of SLE. Across most clinical phenotypes, type I interferon signaling was upregulated, as well as the JAK-STAT pathways related to IL2 and IL6 signaling, complement cascade, and TNF signaling (Figure 4D). In particular, the IL6/JAK/STAT3 pathway was prominently upregulated in acute cutaneous lupus (malar rash), discoid lupus, and SCLE. They also exhibited strong and significant upregulation of complement pathways and the IL6/JAK-STAT3 pathways alongside the interferon pathways. Significant activation of the IL2/STAT5 pathway, which also involves JAK1/JAK3, were also seen in malar rash, SCLE, and lupus panniculitis. This suggests a potential supplemental role for JAK signaling in a subset of SLE patients with SCLE and discoid lupus. The CD19 B cell compartment also showed a high degree of metabolic activity and cell turnover, suggesting high proliferation rate. As expected, photosensitive conditions such as SCLE, malar rash, and photosensitivity showed upregulation of the UV response pathways. Hierarchical clustering identified three sub-clusters of clinical phenotypes within the CD19 compartment. In particular, the pathway signatures were similar between alopecia, photosensitivity, malar rash, and mucosal ulcers. Separately, discoid lupus, SCLE, and lupus panniculitis clustered together. In contrast, livedo reticularis and skin ulcers exhibit fewer upregulated inflammatory pathways.

In the NK cell compartment, we observed fewer differentially expressed genes and fewer significantly enriched pathways overall. However, upregulated NK cell pathways appear to play a role in alopecia, skin and mucosal ulcers, discoid lupus, and livedo, but to a lesser degree in malar rash or photosensitivity (Figure 4E). From hierarchical clustering, we identify mucosal ulcers and alopecia with similar pathway signatures in NK cells, with upregulation of pathways related to cellular metabolism, turnover, proliferation, as well as type I interferon and TNF signaling. Unlike in CD4, CD14, or B cells, NK cells do not experience as significant upregulation of interferon or JAK/STAT pathways. Interestingly, many inflammatory pathways in NK cells are downregulated in SCLE and lupus panniculitis, suggesting a potential regulatory role of NK cells in these subtypes, and that downregulation of inflammatory pathways in NK cells could be related to these phenotypes. NK cells exhibit few differentially upregulated pathways for patients with malar rash and photosensitivity.

### Cell-type enrichment in cutaneous disease states in SLE

Finally, after defining cell-type specific and globally dysregulated pathway fingerprints for each lupus-related rash, we then asked the question whether we could detect significant relative differences in the cell-type enrichment in the blood of patients with specific lupus-related rashes. We conducted cell type enrichment analysis using XCell (28) on the bulk transcriptome of whole blood from the same cohort. We identified a total of 17 immune cell types in the blood that had significant enrichment in the SLE cohort, including CD4 naïve and memory T cells, naïve and memory B cells, naïve and memory CD8+ T cells, monocytes, neutrophils, and plasma cells. The strongest normalized cell type enrichments signatures were seen in the CD4+ memory and naive T cells, naïve and memory CD8+ T cells, monocytes, B cells (Figure 5A). Larger positive values indicate a higher enrichment score. For several rashes, we identified statistically significant changes (padj < 0.05) in the cell type enrichment in the blood, including discoid lupus, mucosal ulcers, alopecia, and in photosensitivity (Figure 5B-E). There was an increase in class-switched memory B cells and in plasma cells in patients with mucosal ulcers (Figure 5C). In alopecia and in photosensitivity, there was a decrease in relative enrichment score in CD8+ naive T cells, suggesting an immunoregulatory role for this cell type in those rashes. In contrast, there was a corresponding increase in enrichment of CD4+ T cells for alopecia and photosensitivity (Figure 5B, D). In discoid lupus, there was strong relative increases in enrichment score for B cells in the blood, including both naive and memory B cells (Figure 5E). All other comparisons of cell type enrichment in the global blood compartment were not significant.

**Figure 5:**
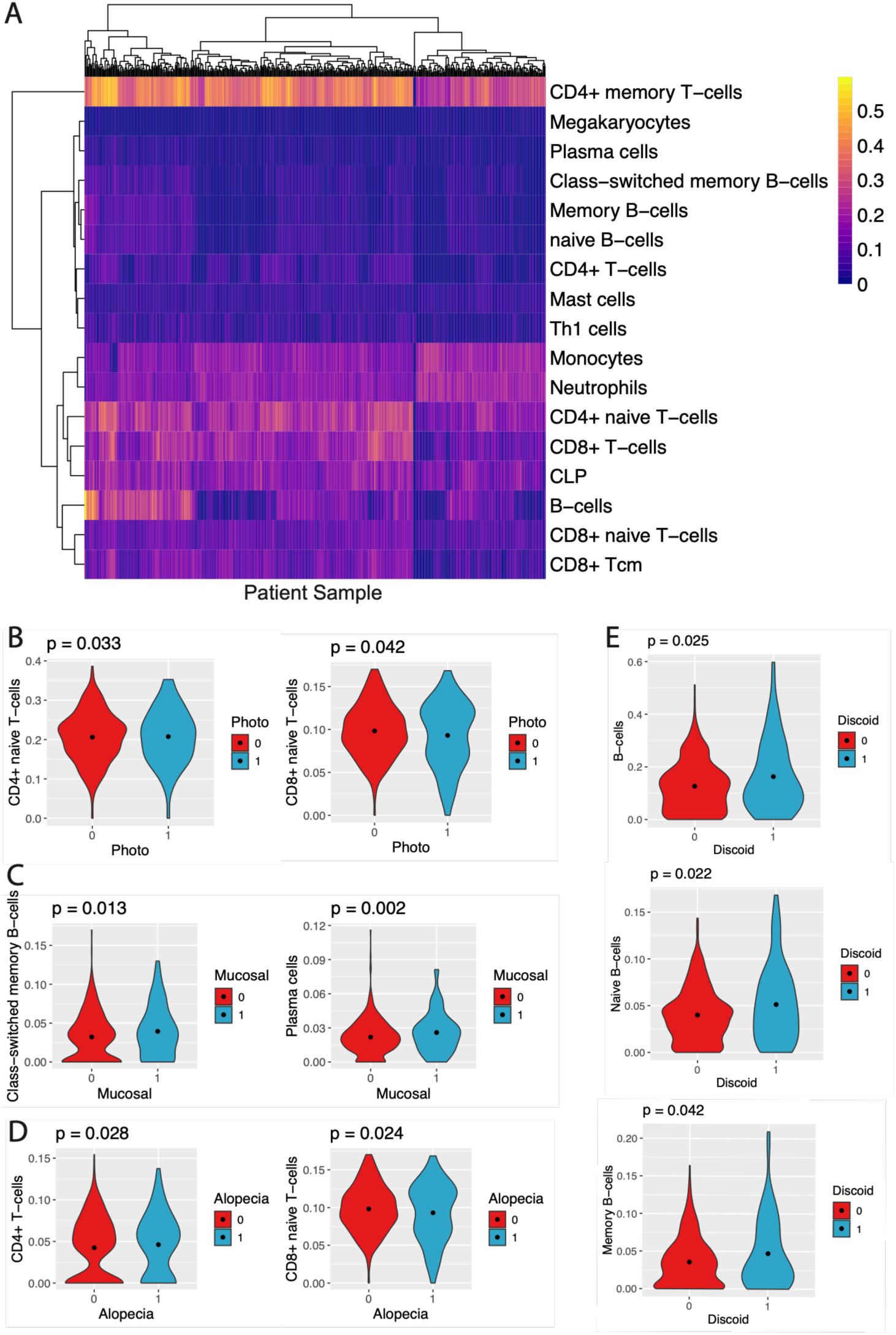
Cell type enrichment in cutaneous disease states in SLE. (A) Cell type enrichment calculated with XCell from bulk RNAseq from all CLUES patients (N = 276), with normalized abundances for each predicted cell type. Significant differences (padj < 0.05) were detected in enrichment scores for CD4+ and CD8+ T cells for patients with photosensitivity (B), in memory B cells and plasma cells for patients with mucosal ulcers (C), and in CD4+ and naive CD8+ T cells for patients with alopecia (D). Significant differences were also observed in enrichment of B cells including naive and memory B cells in discoid lupus (E).

## Discussion

This study provides a comprehensive, large-scale transcriptomic analysis of cutaneous phenotypes from a longitudinal cohort of systemic lupus erythematosus (SLE) patients, allowing us to identify key molecular pathways involved in specific skin-related manifestations of lupus. By utilizing both whole blood bulk RNA sequencing and immune cell-sorted RNA sequencing data from the CLUES cohort, we have successfully mapped unique molecular correlates associated with 10 different rashes observed in SLE patients. We conducted parallel analyses of bulk RNAseq and cell-sorted RNAseq data, which allowed for a more refined understanding of immune cell contributions to cutaneous manifestations in SLE. The integration of these datasets allowed us to construct an interactome network by overlapping differentially expressed genes across various cutaneous manifestations of SLE. This network analysis highlighted shared pathways between rash subtypes, contributing to a deeper understanding of the molecular underpinnings of these manifestations. Hallmark pathway analysis of both unsorted and cell-sorted data identified distinct molecular pathway “fingerprints” for different rashes, providing critical insight into the immune and molecular mechanisms underlying these skin findings. Cell-type-specific findings further illuminate the cellular contributors to specific cutaneous phenotypes. Our results have direct therapeutic implications for patients that suffer from specific cutaneous phenotypes of SLE.

One key unexpected finding from our analysis was that the type I interferon (IFN-α) pathway, known as a defining hallmark of lupus disease activity and pathogenesis, is not universally upregulated across all cutaneous subtypes, suggesting that targeting IFN-α may be more effective in certain lupus subtypes compared to others. We found that IFN-α and IFN-gamma pathways were upregulated in patients with acute cutaneous lupus (malar rash), discoid lupus, SCLE, alopecia, livedo reticularis, and lupus panniculitis. In contrast, patients with skin ulcers and mucosal ulcers did not have significant upregulation of interferon pathways. This absence of IFN signaling in certain cutaneous phenotypes suggests that SLE skin manifestations may involve different immunological pathways, underscoring the heterogeneity of the disease. This also represents a potential targetable therapeutic opportunity for patients with refractory lupus cutaneous phenotypes that tend to have stronger type I IFN signatures. This suggests that anifrolumab (Saphnelo), a biologic medication targeting IFN-α which is approved in the US for moderate to severe SLE, is a potential treatment for refractory cutaneous lupus in patients with SLE.

Subacute cutaneous lupus erythematosus is a cutaneous subtype of lupus that is also poorly understood. Our findings show that patients with SCLE exhibited a broad and strong upregulation of several cytokine immune pathways, including IFN-α, IFN-γ, TNF-α, IL2-STAT5 signaling, and IL6-JAK-STAT3 signaling particularly in B cells and CD4 T cells. The involvement of both IFN and multiple JAK-related pathways in specific immune cell types suggests that SCLE patients with SLE may benefit from treatment with JAK inhibitors as well as type I interferon blockade. The other cutaneous phenotypes that had strong upregulation of JAK pathways were in discoid lupus erythematosus and acute cutaneous lupus. These findings is particularly noteworthy given the emerging role of JAK inhibitors in autoimmune diseases. Several FDA approved JAK inhibitors include topical (ruxolitinib) and systemic (upadacitinib and abrocitinib) formulations, which are being used for vitiligo and atopic dermatitis, respectively. Our findings provide a direct rationale for investigating either systemic or topical JAK inhibitors as a potential treatment for SCLE and discoid lupus in a real-world clinical trial.

CD4 T cells are known to play a central in the pathogenesis of SLE (29, 30), but the role of specific pathways dysregulated in the blood compartment related to skin manifestations are not well studied. From our cell-type specific analysis, we identified significant enrichment of differentially expressed genes in CD4 T cells and B cells, suggesting a prominent role of these immune cells in the pathophysiology of both SCLE and livedo reticularis. SCLE and livedo reticularis shared many common upregulated genes in the CD4 T cell compartment, further highlighting the importance of T-cell specific responses in these rashes. Similarly, our study identifies specific pathways involved in UV response that are shared by SCLE and photosensitivity, indicating a potential unexpected role of CD14+ monocytes in the pathophysiology of photosensitivity in lupus.

We also identified that alopecia in SLE patients, which is a common and psychosocially distressing phenotype, was associated with upregulation of heme metabolism pathways. Iron deficiency has been known to be linked to hair loss conditions such as androgenetic alopecia. This observation suggests that therapeutic strategies aimed at addressing iron metabolism, especially in patients with anemia, may be beneficial in treating lupus-related alopecia. Surprisingly, mucosal ulceration, alopecia, and livedo reticularis were associated with upregulation of inflammatory programs in NK cells, suggesting that NK cells may play an underappreciated role in the pathogenesis of these cutaneous manifestations. Cell type enrichment analysis further predicted differential enrichment of specific immune cell types in various cutaneous manifestations of SLE, including malar rash, discoid rash, SCLE, alopecia, and photosensitivity. These findings suggest that different immune cells may be preferentially involved in the development of distinct rashes, further supporting the notion of immune heterogeneity in SLE.

A strength of this study is the robust cohort of SLE patients and the availability of high-quality large-scale transcriptomic data, which allows us to obtain significant insights into the molecular basis of lupus-related cutaneous manifestations. The CLUES cohort is one of the largest SLE cohorts in the world with significant genetic and ethnic diversity and rich phenotyping data. Furthermore, parallel use of bulk RNAseq and immune cell-sorted RNAseq data allows for a deeper understanding of the specific immune cells contributing to different rash subtypes. Pathway and interactome analysis identified shared and unique molecular pathways, with potential therapeutic targets, such as JAK inhibitors for SCLE. We also discovered unexpected roles for CD14+ monocytes in photosensitivity and NK cells in alopecia, mucosal ulceration, and livedo reticularis, which identifies new avenues for therapeutic development. A weakness of this study is the use of bulk RNAseq data from whole blood, which may not fully capture the complexity of local immune responses at the site of the skin rash. At the same time, the use of bulk RNAseq allows us to quickly profile several hundred patients at once across a large cohort, since conducting scRNAseq across such as large cohort is very costly. Furthermore, scRNAseq can be biased toward immune cell types that are more abundant in these patients, and are typically only done on PBMCs, ignoring contributions from other important cell types such as the granulocytes of the myeloid compartment. Although the study investigates blood immune signatures across multiple cutaneous subtypes, it does not incorporate skin biopsy samples at the time of blood transcriptomics to directly assess the immune responses within affected tissues, which could provide additional valuable insights. At the same time, the goal of the study was to identify molecular signatures from whole blood that are predictive of rashes without requiring a skin biopsy, which can only be done on active lesional skin. While our findings support potential therapeutic targets and modalities, including JAK inhibitors, further clinical validation and functional studies are required to confirm the efficacy and safety of these treatments in specific subgroups of SLE patients, including those with low disease burdens. The cohort size and demographic characteristics may limit the generalizability of the findings, requiring replication in larger, more diverse populations to confirm the observed molecular pathways and immune cell signatures. One limitation of this study is related to recruitment primarily patients with SLE disease without matched controls. As a result, the objective of this study was to identify molecular markers that could be used to predict cutaneous phenotypes among a cohort with known SLE disease, rather than differentiate SLE patients from controls. Another limitation of this study is that a subset of patients in this cohort had historical manifestations of cutaneous lupus that may or may not be flaring or active at time of clinical sampling. We recognize that all clinical manifestations of lupus are not fixed, and dynamic fluctuation of disease activity is characteristic of autoimmune inflammatory conditions.

Taken together, our conclusions represent a significant step forward in understanding the molecular pathways and immune cell involvement in SLE-related skin manifestations, spanning both common and rare phenotypes. The identification of pathway “fingerprints” and novel therapeutic targets holds promise for the development of more personalized and effective treatments for SLE patients, particularly those with specific cutaneous manifestations, and may allow us to predict which patients may benefit from specific therapies to advance personalized medicine for SLE.

## Materials and Methods

### Study design

Participants were recruited from the California Lupus Epidemiology Study (CLUES) as outlined previously (*15*). CLUES was approved by the Institutional Review Board of the University of California, San Francisco. All participants signed a written informed consent to participate in the study. Study procedures involved an in-person research clinic visit, which included collection and review of medical records prior to the visit; history and physical examination conducted by a physician specializing in lupus; a collection of biospecimens, including peripheral blood for clinical and research purposes; and completion of a structured interview administered by an experienced research assistant. During the interview, comprehensive clinical characteristics were documented including overall disease activity (Systemic Lupus Erythematosus Disease Activity Index score (SLEDAI) and lupus severity index), medications, self-reported race/ethnicity, sex, and specific cutaneous manifestations related to SLE. All SLE diagnoses were confirmed by board-certified study physicians based upon one of the following definitions: (a) meeting ≥4 of the 11 American College of Rheumatology (ACR) revised criteria for the classification of SLE as defined in 1982 and updated in 1997, (b) meeting 3 of the 11 ACR criteria plus a documented rheumatologist’s diagnosis of SLE, or (c) a confirmed diagnosis of lupus nephritis, defined as fulfilling the ACR renal classification criterion (>0.5 grams of proteinuria per day or 3+ protein on urine dipstick analysis) or having evidence of lupus nephritis on kidney biopsy. Included in this data collection were cutaneous manifestations of lupus. In this study, we focused on a subset of 10 different rashes and/or cutaneous manifestations related to SLE: acute cutaneous lupus (malar rash), discoid lupus (DLE), photosensitivity, mucosal ulcers, skin ulcers, alopecia, subacute cutaneous lupus (SCLE), lupus panniculitis, livedo reticularis, and other rashes related to SLE. We focused the analysis on skin phenotypes that appeared in at least 10 patients within the entire cohort. Clinical phenotypes that were excluded from our analysis due to low incidence include bullous lupus erythematosus, calcinosis cutis, lupus perniosis, and sclerodactyly. All clinical data collected at sampling and the self-reported race was used for downstream analyses.

### RNA-seq processing and quality control

The two datasets utilized in this study includes bulk RNA sequencing data from whole blood and cell-sorted peripheral blood mononuclear cells from the CLUES cohort. Whole blood was collected from patients into PAXgene tubes and underwent human rRNA and globin RNA depletion followed by library preparation using the NEBNext library preparation kit. Paired end Illumina sequencing was then carried out on a Novaseq6000. Host gene counts were generated by pseudoalignment against the human transcriptome using Kallisto (31). Adapter-trimmed reads were used as input. We quantified gene expression using raw counts and kept the genes that showed an average FPKM value across the samples >0.5. FastQC (v0.11.8) was run to assess the quality of the sequence reads. For the sequencing on sorted immune cells, samples were isolated from patient donors as outlined previously (16). Cells were isolated from peripheral blood utilizing magnetic beads (CD14+ monocytes, B cells, CD4+ T cells, and NK cells) using EasySep protocol from STEM cell technologies on 120 patients totaling 480 samples. RNA was extracted using a Qiagen column and quality was assessed on Agilent bioanalyzer. Libraries were sequenced on HiSeq4000 PE150. Salmon v0.8.2 (32) was used for our alignment-free pipeline. Adapter-trimmed reads were used as input. We quantified gene expression using raw counts and kept the genes that showed an average FPKM value across the samples >0.5. FastQC (v0.11.8) was run to assess the quality of the sequence reads. Genes with low expression were filtered out before statistical testing. The cell sorted data is deposited and publicly available on the Gene Expression Omnibus (GEO) website under the accession GSE164457. The other data is available on request from the authors.

### Differential expression and pathway analyses

We performed differential expression gene testing with DESeq2 (v.1.24.0 R package) (*19*) using default settings. Sequence lane, medication at time of sampling, sex, batch/plate, and race were entered as covariates within the DESeq2 model using a negative binomial generalized linear model. We conducted differential expression by comparing patients with and without specific cutaneous clinical manifestations of SLE. For example, we conducted differential expression analysis for malar rash using DESeq2 by comparing patients with and without malar rash, with patients without malar rash as the reference baseline. Similarly, we conducted differential expression analysis on the cell-sorted bulk RNAseq dataset (CD4, CD14, CD19, NK cell) in the same manner. Statistical significance was set a 5% FDR (Benjamini-Hochberg). Differential expression analysis was conducted for all 10 cutaneous phenotypes that had at least 10 patients exhibiting the phenotype. For pathway analyses to identify immune pathways and signatures associated with each cutaneous phenotype, we inpuwed the sorted differenyally expressed gene lists defining the log 2 fold change and the adjusted p-values into the Bioconductor package fgsea (v 1.10.1) for gene set enrichment analysis (GSEA) (*20*) with Hallmark pathways taken from the MSigDB (26).

### Cell type enrichment analyses

The XCell package was used to carry out cell type enrichment analyses (28). This was conducted on the merged and normalized bulk RNAseq data for the enyre cohort. XCell performs cell type enrichment analysis from gene expression for 64 immune and stroma cell types, based on transcriptomic signatures from thousands of pure cell types from various sources as outlined previously (28). The existing compendium of cell type signatures were filtered to remove cell types not relevant to this analysis. The false discovery rate was controlled by the Benjamini–Hochberg method.

### Statistics and reproducibility

The statistical analyses for each part of the approach are described above. A threshold of P = 0.05 for false discovery rate and for adjusted P-values was set. The code is publicly available to ensure the reproducibility of our results.

## Acknowledgments

We are grateful to all patients who participated in this study. We would like to thank the administrative and medical team of UCSF Rheumatology clinics. We would like to thank the PREMIER center as well as members of the Sirota Lab for useful discussion.

## Author contributions

All authors were involved in drafting the article or revising it critically for important intellectual content, and all authors approved the final version to be published. EYL and MS had full access to all the data in the study and take responsibility for the integrity of the data and the accuracy of the data analysis.

## Data and materials availability

The data that support the findings of this study are available from the corresponding author upon reasonable request.

