## Supplemental Information for "Transcriptomic analysis reveals immune signatures associated with specific cutaneous manifestations of lupus in systemic lupus erythematosus"

**Figure S1**: CLUES cohort demographic histograms. (A) age, (B) self-reported race/ethnicity, (C) sex, and lupus severity metrics including (D) lupus severity index and (E) SLEDAI Scores.

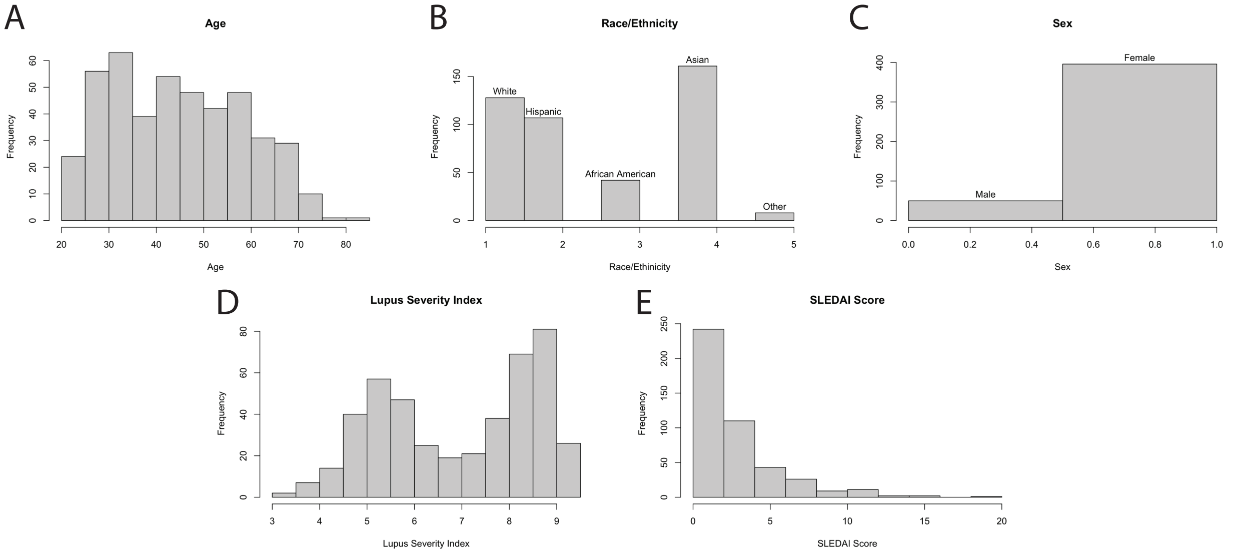

**Figure S2**: Quality control of whole blood bulk RNAseq (A) Histogram distribution of gene counts for all samples. (B) Plot of fraction of counts compared to fraction of genes for all samples. (C) Distribution of gene biotypes (protein coding, pseudogene, lncRNA, or other) for all samples.

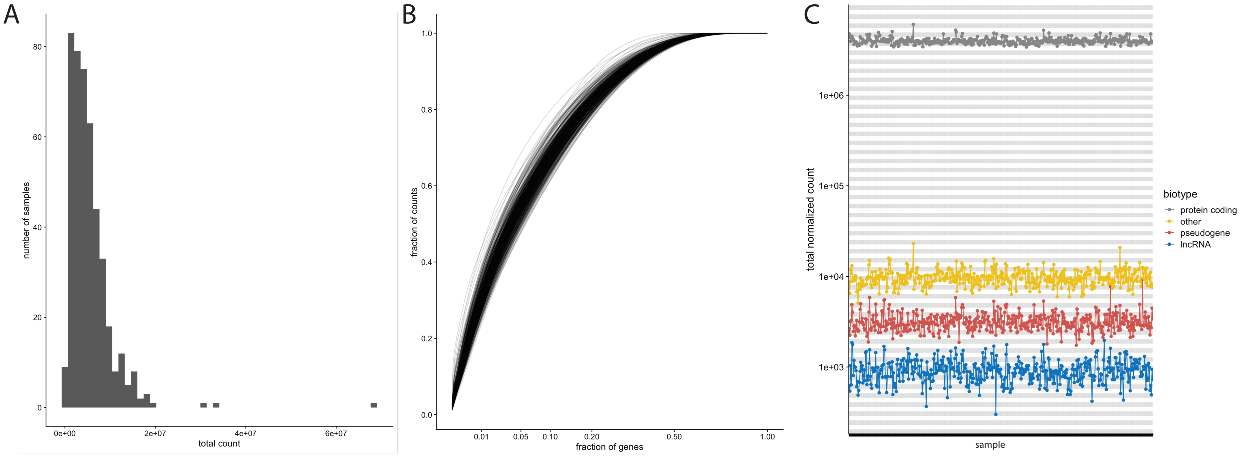

**Figure S3**: Principal component plot of whole blood bulk transcriptome and additional clinical characteristics

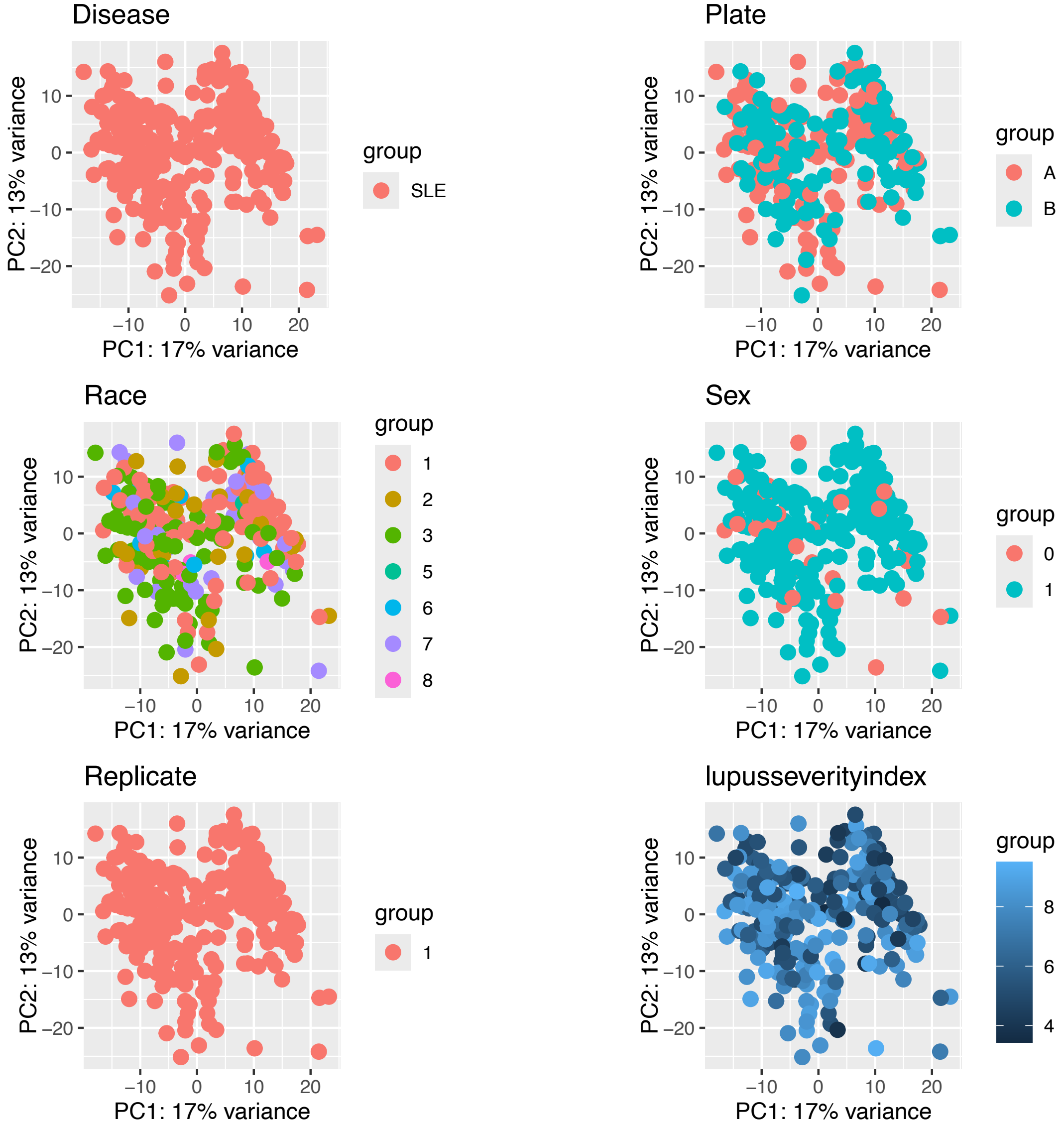

**Figure S4**: Hierarchical clustering of clinical characteristic, highlighting clinical subtypes of SLE-related rashes

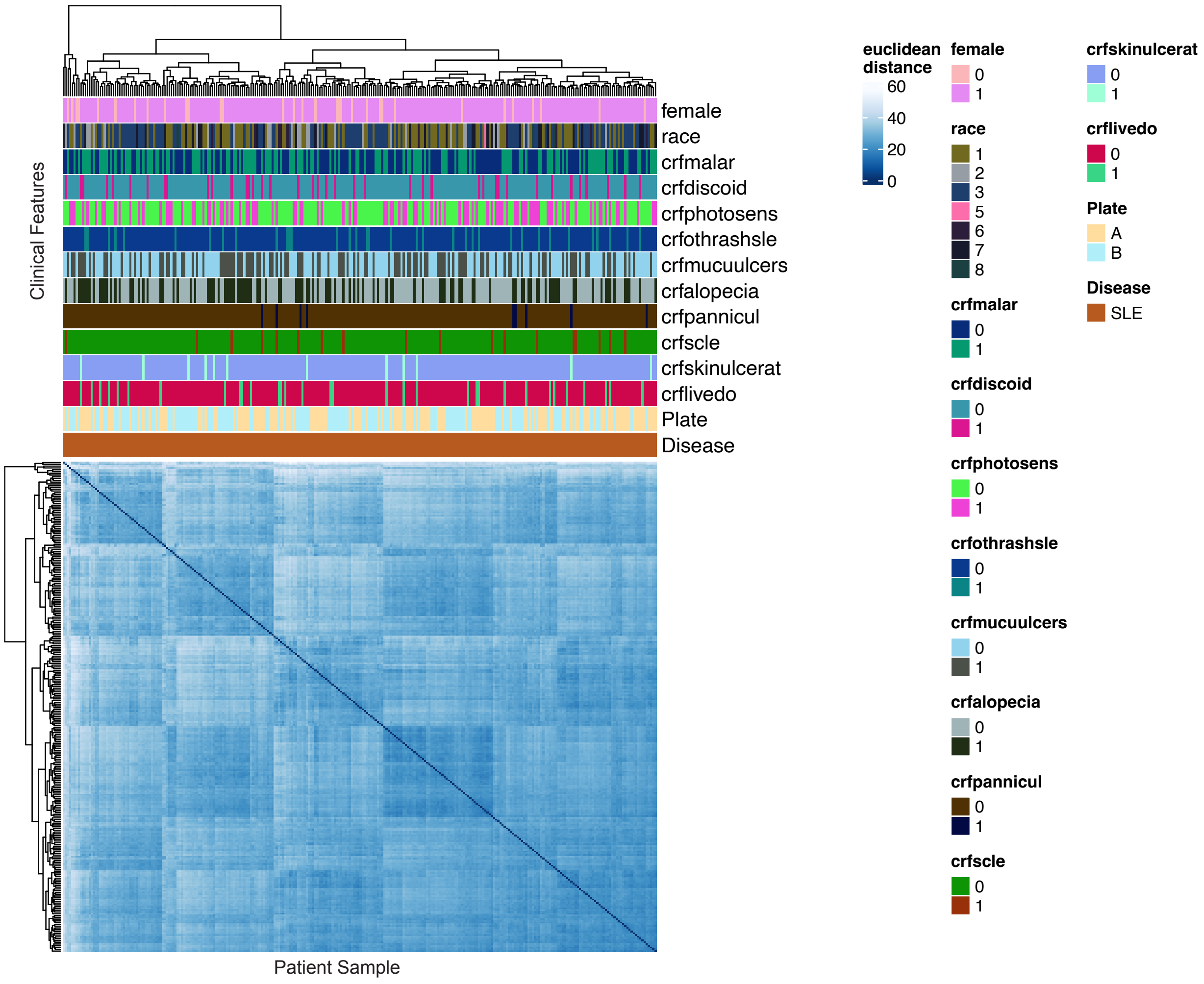

**Figure S5**: Upset plots on overlapping differentially expressed genes for SLE rash subtypes in (A) whole blood RNAseq, and in cell sorted RNAseq data for (B) CD4 T cells, (C) CD14 monocytes, (D) CD19 B cells, and (E) NK cells.

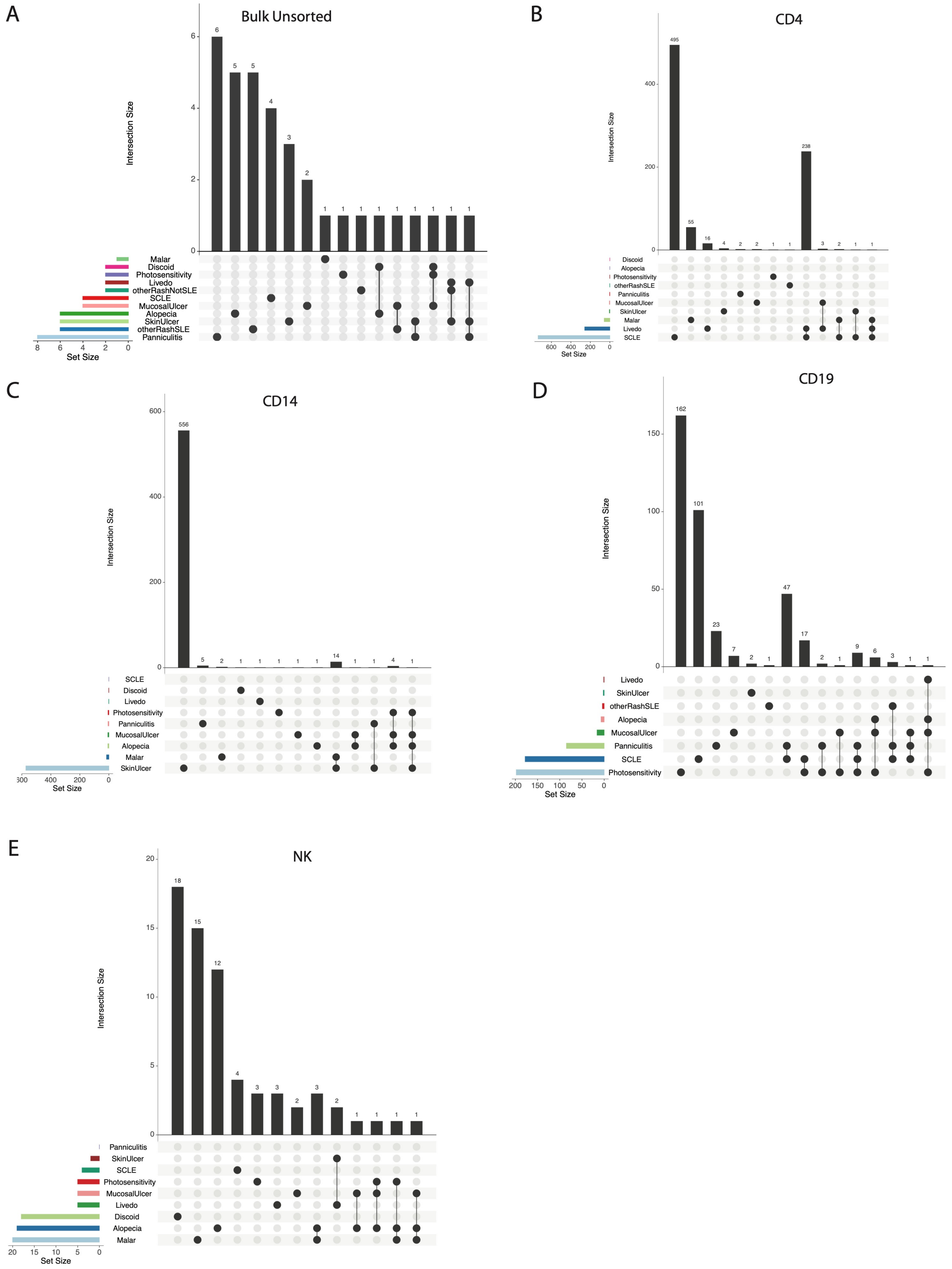

**Figure S6**: Pathway enrichment plots from gene set enrichment analysis of whole blood bulk RNAseq for each SLE rash subtype (A)-(K).

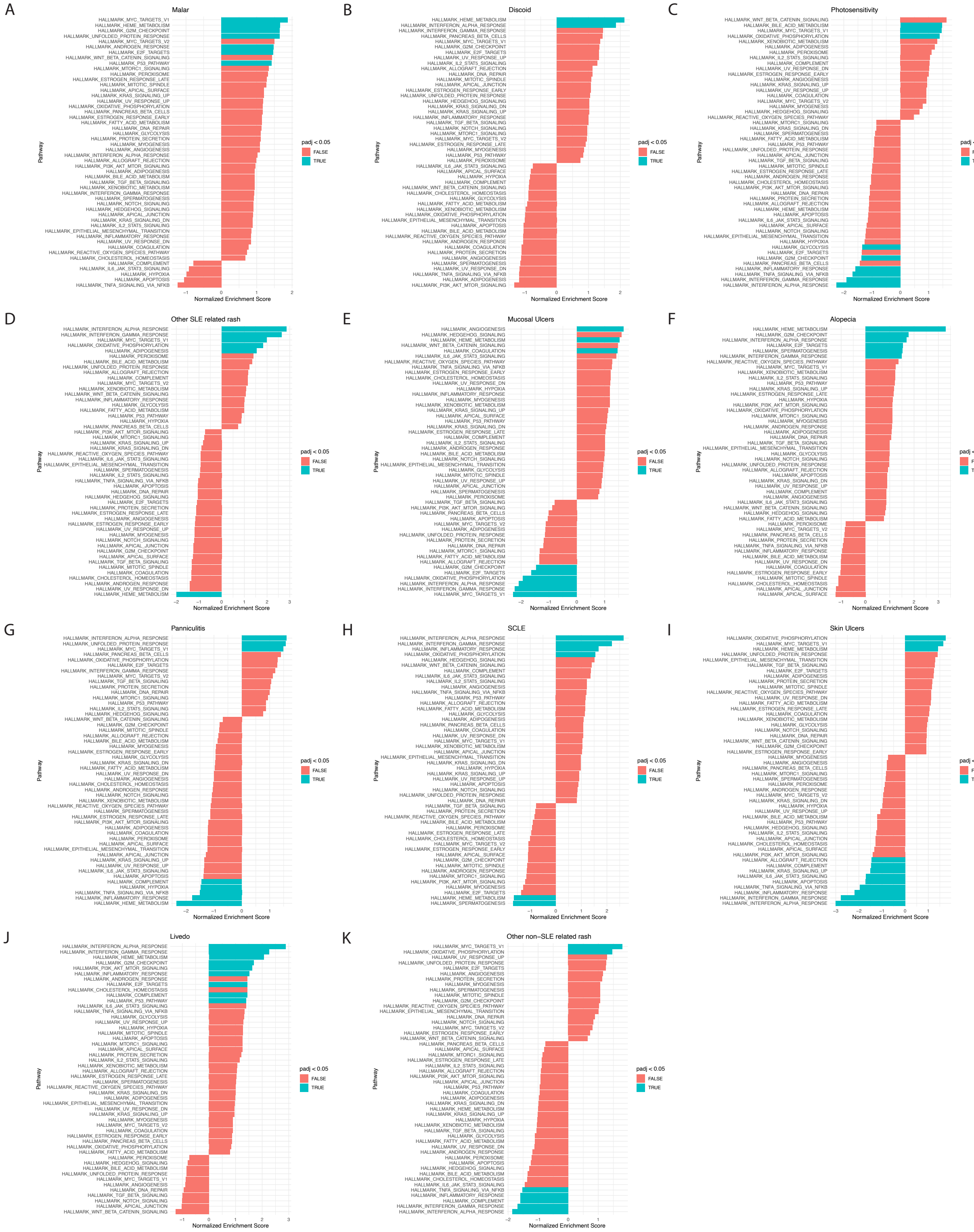
